# Practical selection of representative sets of RNA-seq samples using a hierarchical approach

**DOI:** 10.1101/2021.02.04.429817

**Authors:** Laura H. Tung, Carl Kingsford

**Affiliations:** Computational Biology Department, School of Computer Science, Carnegie Mellon University, Pittsburgh, PA 15213, USA

## Abstract

Despite numerous RNA-seq samples available at large databases, most RNA-seq analysis tools are evaluated on a limited number of RNA-seq samples. This drives a need for methods to select a representative subset from all available RNA-seq samples to facilitate comprehensive, unbiased evaluation of bioinformatics tools. In sequence-based approaches for representative set selection (e.g. a k-mer counting approach that selects a subset based on k-mer similarities between RNA-seq samples), because of the huge number of available RNA-seq samples and the large number of k-mers/sequences in each sample, computing the full similarity matrix between all samples using k-mers/sequences for the entire set of RNA-seq samples in a large database (e.g. the SRA) has memory and runtime challenges, making direct representative set selection infeasible with limited computing resources. Therefore, we developed a novel computational method called “hierarchical representative set selection” to handle this challenge. Hierarchical representative set selection is a divide-and-conquer-like algorithm that breaks the representative set selection into sub-selections and hierarchically selects representative samples through multiple levels. We demonstrate that hierarchical representative set selection can achieve performance close to that of direct representative set selection, while largely reducing the runtime and memory requirements of computing the full similarity matrix (up to 8.4X runtime reduction and 4.7X memory reduction for 10000 samples that could be practically run with direct subset selection). We show that hierarchical representative set selection substantially outperforms random sampling on the entire SRA set of RNA-seq samples, making it a practical solution to representative set selection on large databases such as the SRA.

## 1 Introduction

A vast number of RNA-seq short-read samples are publicly available at large sequence databases (e.g. NIH’s Sequence Read Archive [12], known as SRA). However, most bioinformatics tools for RNA-seq analyses are evaluated on a limited number of samples; this evaluation may be insufficient, as the tools may not be adequately evaluated by samples with a variety of cell/tissue types and disease conditions. To ensure general applicability, an RNA-seq analysis tool should be validated on varying cell/tissue types and experiments. On the other hand, using all available RNA-seq samples to evaluate RNA-seq analysis tools is infeasible, and many samples in databases are similar to each other. This leads to a need to select a representative subset from available RNA-seq samples that effectively summarizes a large collection of RNA-seq samples to capture various essential transcriptional phenomena. Moreover, bioinformatics tools have algorithm parameters to be optimized, and automatic learning of optimal parameters can be made more robust using representative samples. Thus, our objective is to develop a computational method of selecting a representative subset from a large collection of RNA-seq short-read samples for a given organism (e.g. human), such that RNA-seq analysis tools can be effectively evaluated on this subset. Bioinformatics tools such as transcript assemblers, read mappers, and expression abundance estimators would benefit from a good selection of RNA-seq samples in their evaluation and parameter optimization.

Various representative set selection methods that solve the problem of finding a subset of data points (representatives) to efficiently describe the original collection of data have been developed in the fields of computer vision, signal/image processing, information retrieval, and machine learning. Most common applications of representative set selection include image, video, and text summarizations. Machine learning tasks such as classification and regression can also improve in terms of fast training and reduced memory usage by using a representative subset as the training set [6, 9]. One category of representative set selection methods is clustering-based algorithms [5, 8], using k-means, k-medoids, spectral clustering, or DBSCAN (Density-Based Spatial Clustering of Applications with Noise). Another category is sparse modeling-based algorithms [20, 7], which formulate the representative set selection as a dictionary learning problem, based on the assumption that the entire set can be reconstructed by linear combinations of dictionary items. There are also Rank Revealing QR algorithms [2] that use matrix factorization to find a subset of columns of the data matrix corresponding to the best conditioned submatrix, and algorithms using mutual information and relative entropy to search for a representative subset [17].

In the field of RNA-seq analysis, Hie et al. developed a geometric sketching algorithm [10] for single-cell RNA-seq, which summarizes the transcriptomic heterogeneity within a data set using a representative subset of cells to accelerate single-cell analysis. Using a covering algorithm that approximates the original data space as a union of equal-sized boxes, geometric sketching focuses on even coverage of the transcriptional space spanned by the original set, so that rare cell types can be sufficiently sampled and represented. Maintaining a similar density distribution to that of the original set is useful for video/photo summarizations. However, for RNA-seq analyses, even coverage of the transcriptional space is more important in order to represent rare cell types.

A Python package, apricot [18], has been developed for selecting representative subsets using submodular optimization. Based on the “diminishing returns” property, apricot maximizes a monotone submodular function’s value to find a representative subset. Using facility location functions, apricot maximizes the sum of similarities between each sample and its closest representative sample; as a result, the representative set selected by apricot approximately evenly spans the space of the original data, like geometric sketching. While geometric sketching requires knowing samples’ gene expression vectors, apricot can work with the similarity matrix between the samples directly.

A main challenge in representative set selection for RNA-seq samples is that the number of available RNA-seq samples in large databases is huge and each RNA-seq sample containing read sequences takes up substantial disk space; therefore, it is impractical to download all RNA-seq sequences of all the samples available at a large database like the SRA due to limited disk space. To perform representative set selection directly, we need to obtain gene expression vectors of, or distances between, all available SRA samples; however, SRA streaming is also not feasible due to issues with paired-end reads.

Given this challenge, one might attempt to select a representative set without looking at the sequences of each RNA-seq sample and relying instead on each sample’s metadata, for example, using NCBI’s BioSample attributes [1] that affect gene expression levels to predict gene expression distances between RNA-seq samples. However, for most large RNA-seq collections, including the SRA, the metadata is highly incomplete and most samples do not have the needed metadata values for predicting their gene expression distances.

Thus, we use a sequence-based approach for representative set selection of RNA-seq samples. We randomly sample a small subset of reads from each RNA-seq sample to download, such that the subsets of reads from all available RNA-seq samples at the SRA take a reasonable disk space. We count k-mers in the subset of reads of each sample and compute the similarity between k-mer distributions of samples. This approach selects a representative set based on k-mer similarities and thus sequence similarities among RNA-seq samples. Since the number of publicly available RNA-seq samples in the SRA is large (*N* =196523 for human) and the number of k-mers in each sample is large (∼2000000 k-mers), computing the 196523×196523 similarity matrix with k-mers has memory and runtime challenges even using a chunking method for matrix computation [13].

To tackle this challenge, we developed a novel method called “hierarchical representative set selection.” The hierarchical representative set selection is a divide-and-conquer-like algorithm that hierarchically selects representative samples through multiple levels. At each level, samples are divided into smaller chunks, and representative set selection is performed on each chunk with a weighting scheme. The representative samples selected from every chunk are merged into the next level, and the process repeats until the size of the similarity matrix of the merged samples is feasible for the computing resources.

Our results show that the hierarchical representative set selection achieves performance close to that of the direct representative set selection using apricot, while substantially reducing the runtime and memory requirements of computing the full similarity matrix (up to 8.4X runtime reduction and 4.7X memory reduction for 10000 samples that could be practically run with direct subset selection), thus making selecting representative samples from the entire SRA RNA-seq samples feasible (the estimated runtime reduction is 90X and memory reduction is 41.4X for the SRA full set of 196523 human samples). We demonstrate that the representative subset selected by our hierarchical representative set selection method from all available human RNA-seq samples in the SRA better represents the transcriptomic heterogeneity among those SRA samples than that by random sampling, and thus can be used for more comprehensive and complete evaluation of bioinformatics tools.

## 2 Methods

### 2.1 Problem formulation

Let set *R* be a large set of RNA-seq samples (such as all the RNA-seq samples in the NIH SRA database for a given organism), let *d*(*i, j*) be a distance or dissimilarity measure between samples *i* and *j* in *R*, and let *d*(*x, S*) be the distance between a data point *x* and its closest data point in a set *S*. A reasonable formulation is to find a representative subset 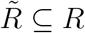, such that is as small as possible.

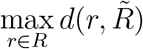

is as small as possible.

This is equivalent to minimizing the classical Hausdorff distance which is defined as: 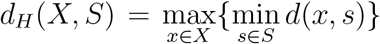 where *X* is the full set and *S* is its representative subset as proposed in [10]. The Hausdorff distance can be used to evaluate how well a selected subset represents the original full set (a smaller value is better) [10]. However, the classical Hausdorff distance is highly sensitive to extreme outliers [11, 19]. Thus, in practice, a more robust measure, the partial Hausdorff distance, is used to evaluate the representative subset [11, 10]. The partial Hausdorff distance is defined as: 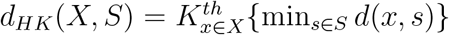 where 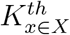 is the *K*^*th*^ largest value (counting from the minimum), and a parameter *q* = 1 − *K*/|*X*| is used to determine *K* (when *q* = 0, *d*_*HK*_ = *d*_*H*_; when *q* is small enough, *d*_*HK*_ is very close to *d*_*H*_ but is robust to extreme outliers) [11, 10].

### 2.2 K-mer similarity-based approach

The similarity between k-mer distributions of RNA-seq samples reflects the similarity between their sequences and is a reasonable approach for computing *d*(*i, j*). Thus, by counting the k-mers of RNA-seq reads, we can select a representative set based on the k-mer similarities and therefore sequence similarities among samples. Downloading all reads of all SRA samples is infeasible, so we download a small subset of reads from each RNA-seq sample. To represent a full RNA-seq sample, the sampled small subset of reads are random with respect to the genome coordinates. For approximately 88% of SRA RNA-seq samples, the reads are stored from the sequencer without alignment. Reads coming from Illumina sequencers without alignment are random with respect to the genome coordinates. Thus, a range of reads downloaded from unaligned samples by *fastq-dump* [15] are random. We download 10,000 reads to represent each unaligned RNA-seq sample. Choosing a proper k-mer size is important, as smaller k-mers give less information about sequence similarities, while larger k-mers may result in fewer matches due to sequencing errors. To select an optimal k-mer size, we plot the number of distinct k-mers with the varying k-mer size for a range of typical read lengths of Illumina (Figs. S1–S5). In the linearly increasing part of the curve, short k-mers match randomly; from the beginning of the horizontal part of the curve, k-mers start to reveal the genome structure. Thus, we want the smallest k-mer size in the horizontal part of the curve, so that the k-mer matching moves from being random to being representative of the read content and is still resilient to sequencing errors. Among the optimal k-mer sizes we obtained for long, medium, and short read-lengths, we choose a compromise 17 as the optimal k-mer size.

Jellyfish [14] was used to count k-mers in the subset of reads. We use canonical k-mers, so all samples are compared based on the common k-mer sequences regardless of the sequenced strand. We use the cosine similarity as the similarity of k-mer distributions between samples. Cosine similarity is commonly used in document clustering and information retrieval; our k-mer vectors have some similar aspects to TFIDF (Term Frequency–Inverse Document Frequency) vectors (e.g. large vocabulary, word counts, high dimension, and high sparsity). Cosine similarity is also a good measure for k-mer-based metagenome comparisons [4].

### 2.3 Hierarchical representative set selection algorithm

To handle the memory and runtime challenges, hierarchical representative set selection (as shown in Algorithm 1) breaks the whole representative set selection into multiple levels of progressive sub-selections, like divide-and-conquer. At each level, the “full set” of samples are divided into smaller equal-size chunks using a seeded-chunking method (see Algorithm 2), such that chunks are well separated with closer samples going into the same chunk if possible. The similarity matrix for each chunk is computed, and weights that determine the size of the representative set for every chunk are computed based on the density/sparseness of each chunk (samples in a denser chunk are more similar to each other). Representative set selection is then performed on each chunk (for example, using apricot’s facility location approach). The representative samples selected from every chunk are merged into a new set, which becomes the “full set” of the next level. This process iterates until the size of the similarity matrix of the merged set is feasible for the computing resources. Lastly, the similarity matrix of the final merged set is computed, and representative set selection is performed on it to get the final representative set of a desired size.

#### Algorithm 1: Hierarchical Representative Set Selection

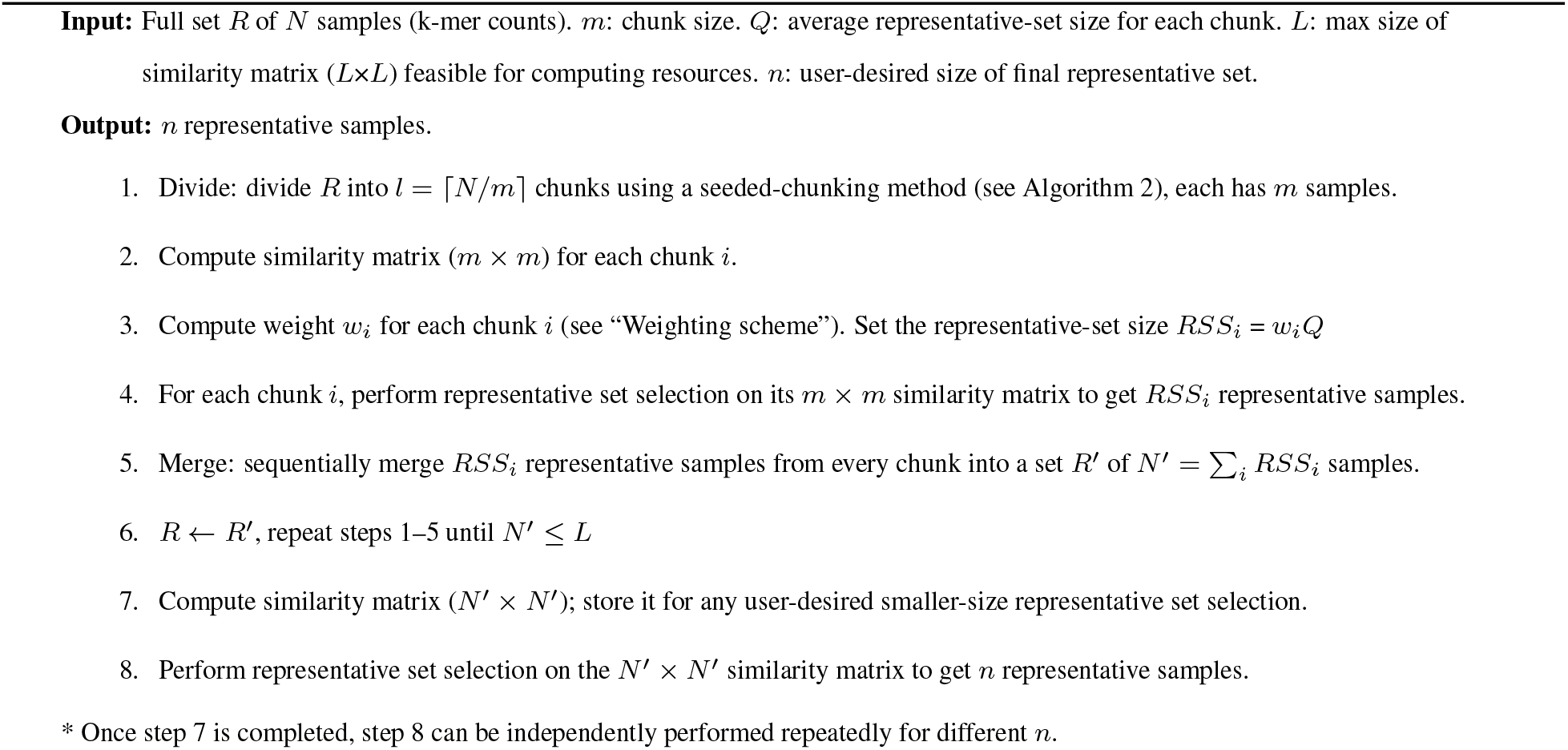

Computing the similarity matrix with k-mers is the runtime and memory bottleneck, while apricot’s runtime is relatively negligible in comparison. For direct representative set selection using apricot, the computational cost for the similarity matrix of the full set is *O*(*N* ^2^). Using the hierarchical selection (considering one iteration of divide-and-merge with *l* chunks, chunk size *m*, and the final merged set size *N*′), the computational cost is reduced to *O*(*lm*^2^)+*O*(*N*′^2^) = *O*(*N*^2^/*l*)+*O*(*N*′^2^). The seeded-chunking has an added computational cost *O*(*Nl*). So the total computational cost of the hierarchical selection is *O*(*N*^2^/*l*) + *O*(*N*′^2^) + *O*(*Nl*), where *l* ≪ *N* and *N*′ ≪ *N*. With multiple iterations, the computational cost is further reduced. Since *m* ≪ *N*, the memory requirement for computing the similarity matrix is greatly reduced.

Fig. 1 illustrates applying the hierarchical representative set selection to 196523 human RNA-seq samples in the SRA, using two levels (two iterations) of divide-and-merge. The first level has *l* = 197 chunks; the second level has *l* = 40 chunks. ∼200 representative samples are selected from each chunk of 1000 samples.

**Figure 1.**
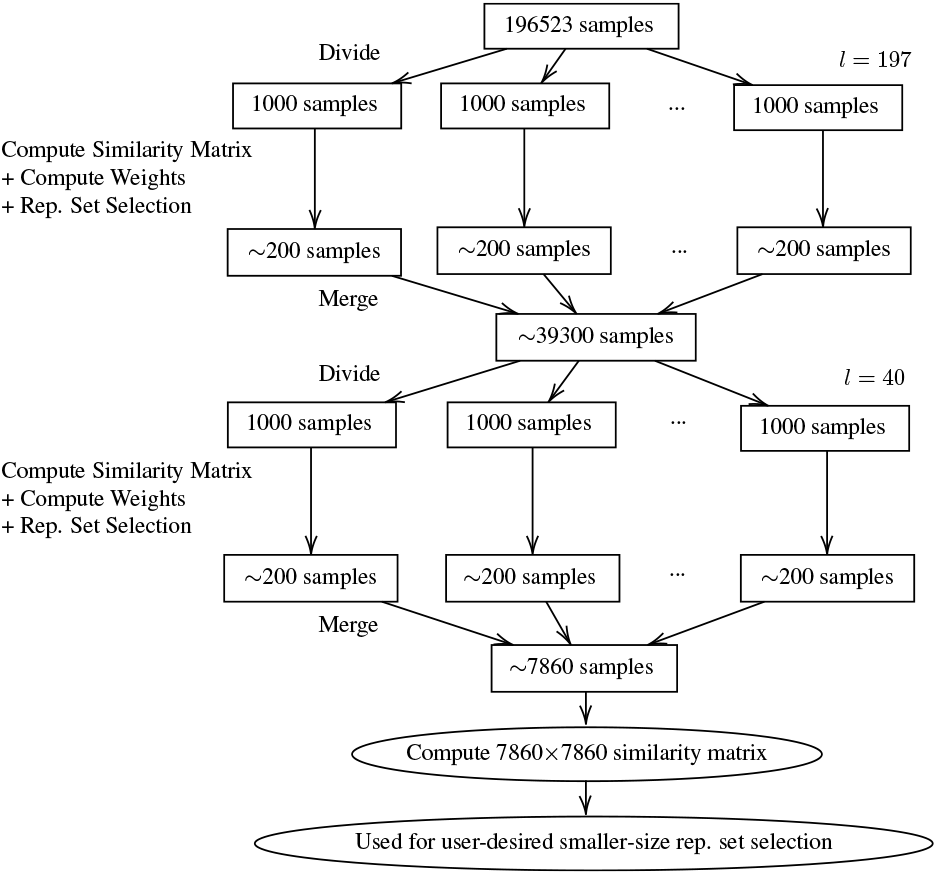
Illustration of the hierarchical representative set selection for 196523 human RNA-seq samples in the SRA.

#### 2.3.1 Seeded-chunking method

The ideal situation for the hierarchical representative set selection is that chunks have no overlaps. Here, the “overlap” of chunks means data points and their close neighbors are in different chunks (e.g. different chunks occupy the same dense cluster). When chunks have no overlaps, the union of the representative sets selected from every chunk would be similar to the representative set selected directly from the original set. When chunks have overlapping regions with similar densities, unnecessarily more representative data points may be selected in the overlapping regions from different chunks. The subsequent representative set selection on the merged set can alleviate this effect, but the over-use of the quota (i.e. a region’s proportion of the desired number of representative data points) in the overlapping regions may cause other regions to have less quota. Thus, more separated chunks lead to more accurate hierarchical selection.

A sequential-chunking method that divides chunks sequentially along the SRA accession list can cause many overlaps, even complete overlaps between chunks. To overcome the introduction of overlaps with sequential chunking, we developed a seeded-chunking method (as shown in Algorithm 2). The seeded-chunking first uses the “farthest point sampling” algorithm [16, 3] to find *l* seeds for *l* chunks, such that the seeds are farthest away from each other. That is, starting from a randomly selected seed, the seeds are chosen one at a time, such that each new seed has the largest distance to the set of already selected seeds (i.e. the largest minimum distance to the already selected seeds). The entire set of RNA-seq samples available at a large database cannot fit into memory at once, but if loading each sample one at a time when computing its distance to a seed, each sample would be repeatedly loaded for *l* − 1 times. Thus, we randomly select a subset *X* from the full set and perform the “farthest point sampling” on *X* to find *l* seeds. Each sample in the full set is assigned to its closest seed. Since chunks have equal sizes, the sample is assigned to its closest seed among all currently non-full chunks (i.e. their current size < *m*).

##### Algorithm 2: Seeded-Chunking Method

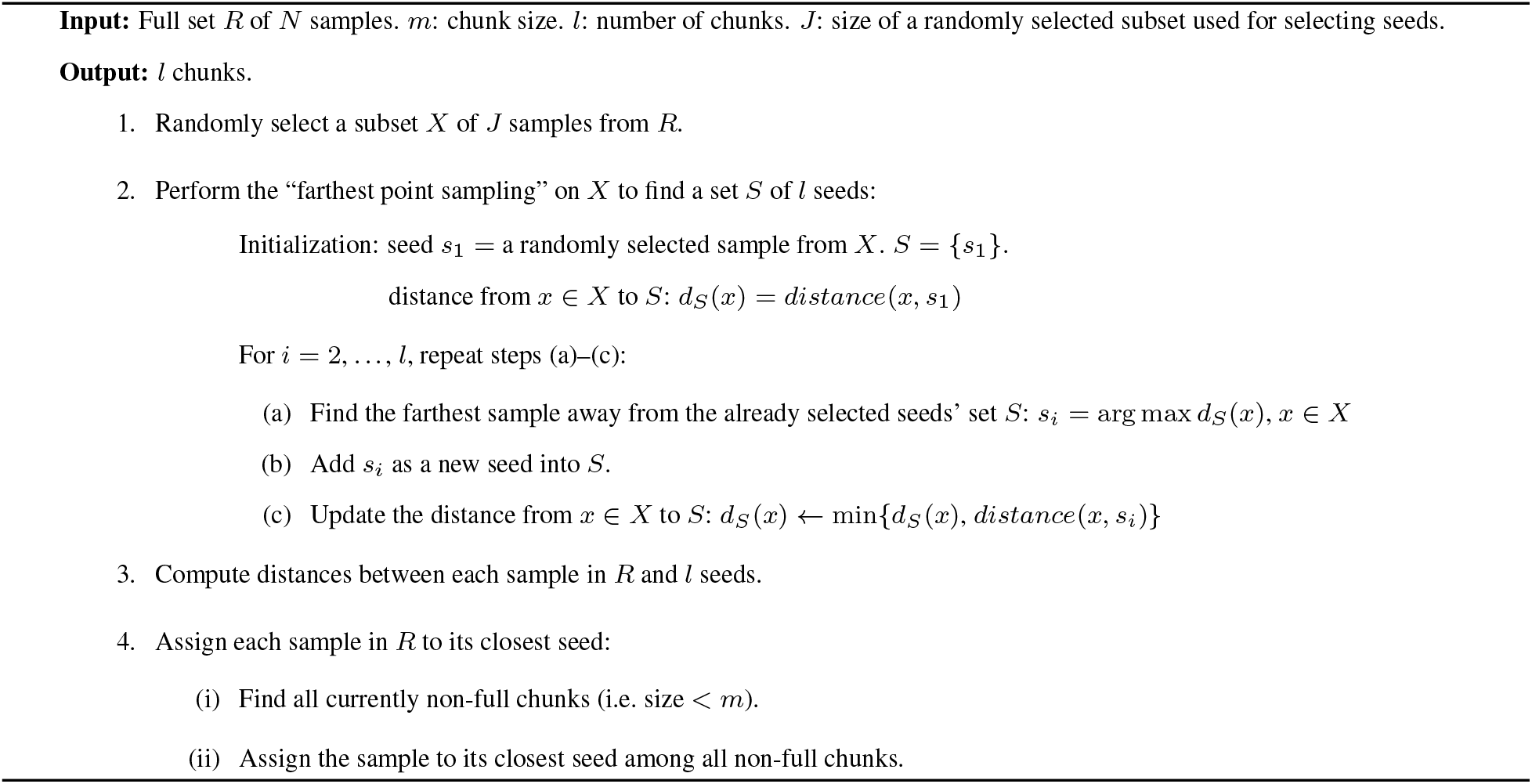

The seeded-chunking can generate well separated chunks, with an added computational cost: *O*(*Nl*) of computing similarities. Since *l* ≪ *N*, this cost is fairly small compared to computing the full similarity matrix (*O*(*N* ^2^)). We benchmarked the seeded-chunking vs. sequential chunking on various sizes of full sets (Table S5). In all cases, the seeded-chunking outperforms the sequential chunking; as the full-set size increases, the seeded-chunking shows more advantages over the sequential chunking in the partial Hausdorff distance.

#### 2.3.2 Weighting scheme

Denser chunks (in which samples are more similar to each other) should have fewer representative samples than sparser chunks, since we want representative data points to approximately evenly span the space of the original data, so that rare cell/tissue types can be sufficiently represented. Since chunks have equal sizes, denser chunks occupy smaller spaces than sparser chunks. Thus, we propose a weighting scheme (“mean^2^-weighting scheme”) to assign the representative-set size to each chunk based on their average density/sparseness.

The mean^2^-weighting scheme is as follows. Let *µ*_*i*_ be the mean of distances between samples in chunk *i*; *z*_*i*_ is the size of chunk *i* (note that when *N*/*m* is not an integer, not all chunks are full); *Q, l*, and *m* are as defined in Algorithm 1. Suppose the *l*th chunk is non-full: let *α*_*l*_ = *z*_*l*_/*m*. The weight *w*_*i*_ and the representative-set size *RSS*_*i*_ for chunk *i* are defined to be:

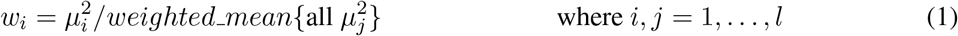

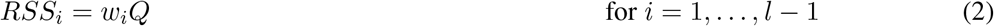

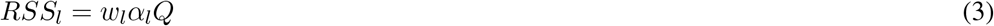

where the 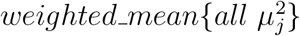 uses *weight* = 1 for *j* = 1, …, *l* − 1 and *weight* = *α*_*l*_ for *j* = *l*. The following relationship holds:

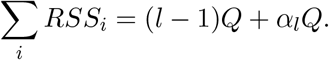

With the seeded-chunking, there could be multiple non-full chunks, but the same weighting method applies. The mean^2^-weighting scheme is a heuristic to adjust the representative-set size for each chunk according to their average density/sparseness.

The code for the hierarchical representative set selection is available at two URLs (the 2nd URL contains the binaries that can also be used for other k-mer similarity applications): https://github.com/Kingsford-Group/hierrepsetselection; https://github.com/Kingsford-Group/jellyfishsim.

## 3 Results

### 3.1 Hierarchical representative set selection achieves performance close to that of direct representative set selection

Excluding SRA samples with no public access permission, aligned samples, and samples with no valid 17-mers (read-lengths < the k-mer size 17, or reads that contain many *N* ‘s in the middle), we obtained 196523 human bulk RNA-seq (Illumina) samples as the SRA entire set. Each sample corresponds to an SRA Experiment.

In this particular implementation of the hierarchical representative set selection, we use apricot’s facility location approach as the base level to perform representative set selection on each chunk and on the merged set. We use cosine distance as the distance measure.

We refer to the direct representative set selection using apricot as “direct apricot”, which includes two parts: (a) computing the similarity matrix of the full set; (b) applying apricot’s facility location approach to the full similarity matrix. The main computational cost of direct apricot comes from part (a).

The motivation of the hierarchical representative set selection is to reduce the runtime and memory requirement of direct representative set selection, while not sacrificing too much accuracy. Thus, we compared the performance (evaluated by *d*_*HK*_) between direct apricot, the hierarchical selection, and random sampling, using the most recent 1000, 2000, 5000, 8000, and 10000 samples in the SRA as the full sets (Fig. 2, Table S1). We also compared the three methods using the early-time 1000, 2000, 5000, 8000, 10000 samples in the SRA as the full sets (in the earliest quarter of the SRA time span, i.e. the 4th quarter of the SRA accession list) (Fig. 3, Table S2), and using the mid-time 1000, 2000, 5000, 8000, 10000 samples in the SRA as the full sets (around the middle of the SRA time span) (Fig. 4, Table S3). For all these cases: we use 1 iteration, *rep*_*set*_*size*/*N* = 0.1, *m* = *N*/*l, Q* = *m*/5. We set *l* = 10, except for *N* = 1000, *l* = 5.

**Figure 2.**
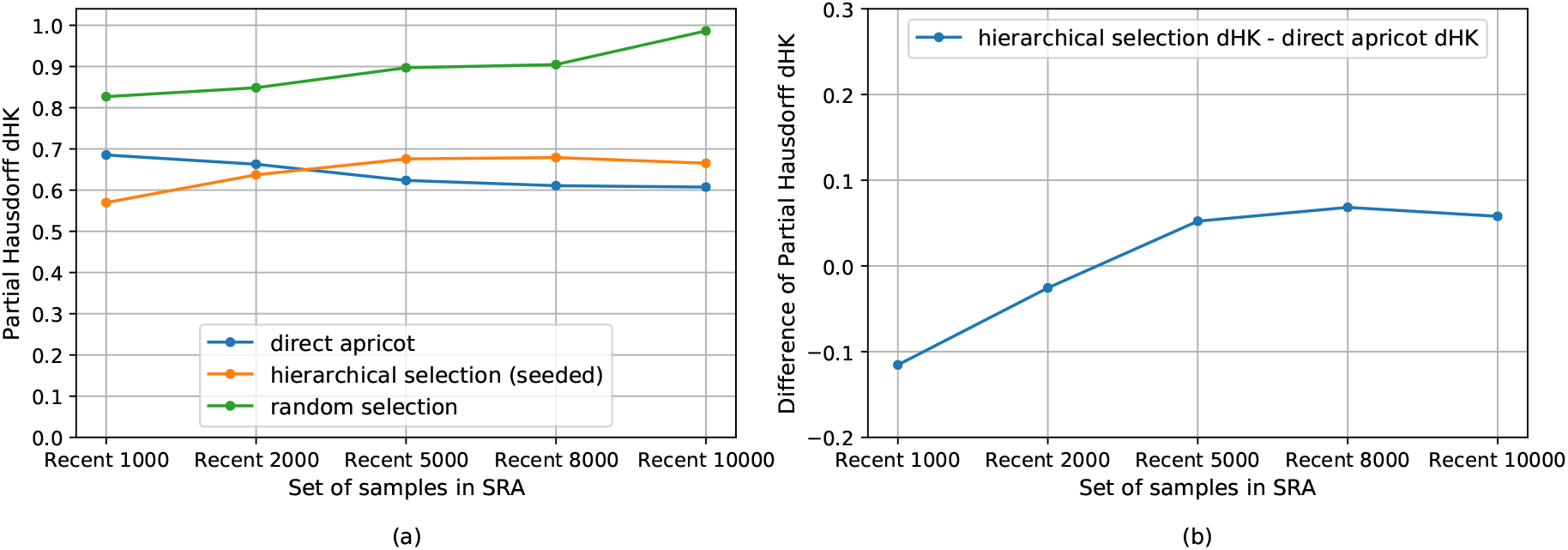
Using the most recent 1000, 2000, 5000, 8000, 10000 samples in the SRA as the full sets. rep set size/N =0.1. (a) Partial Hausdorff distances d_HK_ of direct apricot, hierarchical selection, and random selection. (b) Partial Hausdorff distances’ difference: hierarchical selection d_HK_ − direct apricot d_HK_.

**Figure 3.**
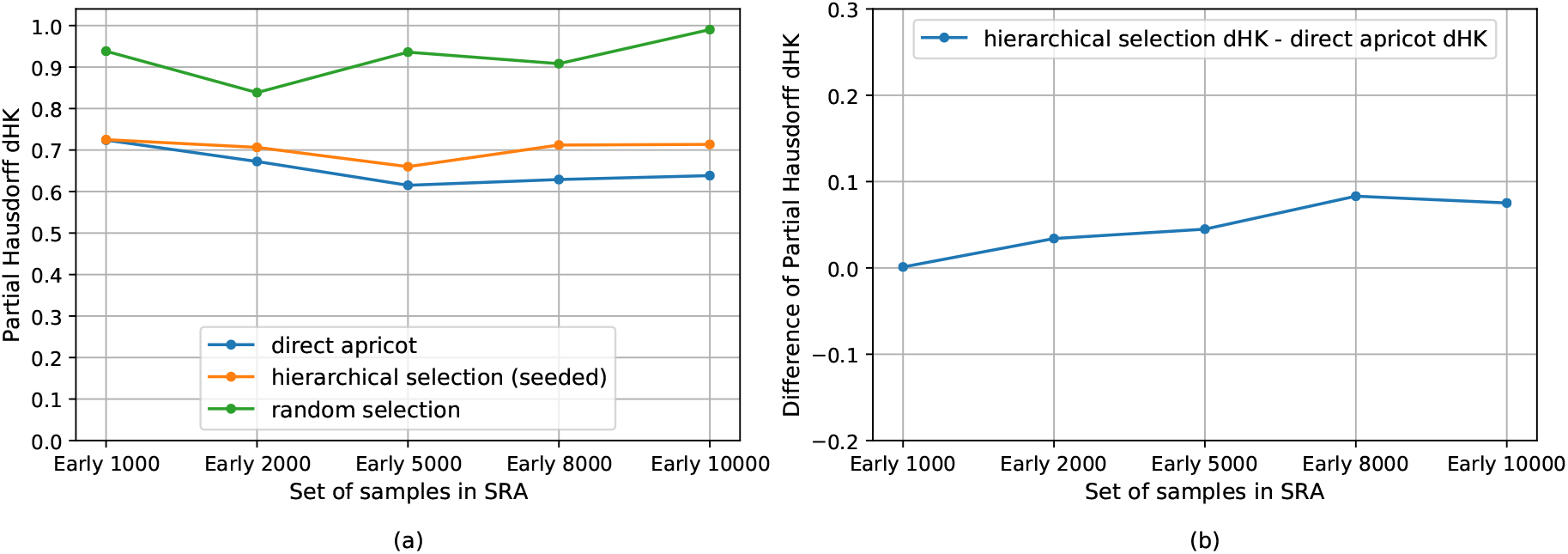
Using the early-time 1000, 2000, 5000, 8000, 10000 samples in the SRA as the full sets. rep set size/N =0.1. (a) Partial Hausdorff distances d_HK_ of direct apricot, hierarchical selection, and random selection. (b) Partial Hausdorff distances’ difference: hierarchical selection d_HK_ − direct apricot d_HK_.

**Figure 4.**
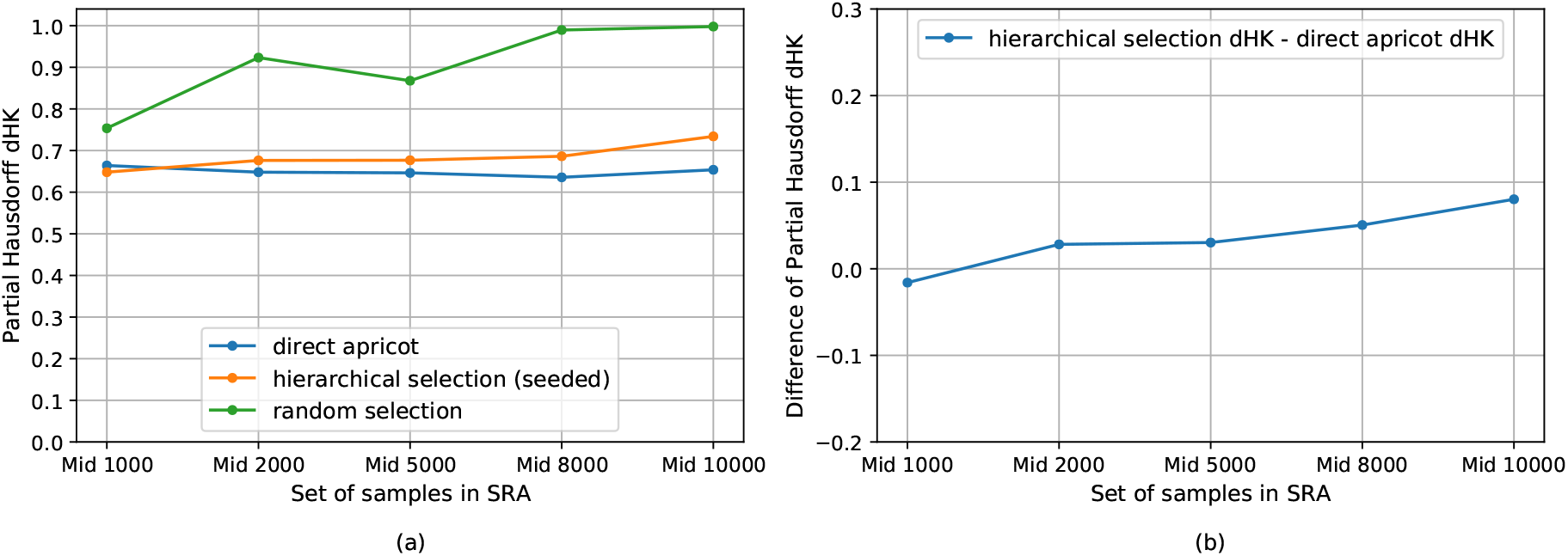
Using the mid-time 1000, 2000, 5000, 8000, 10000 samples in the SRA as the full sets. rep set size/N =0.1. (a) Partial Hausdorff distances d_HK_ of direct apricot, hierarchical selection, and random selection. (b) Partial Hausdorff distances’ difference: hierarchical selection d_HK_ − direct apricot d_HK_.

The hierarchical selection achieves performance close to that of direct apricot. For the recent 1000 and 2000 samples, the hierarchical selection performs better than direct apricot; for the recent 5000, 8000, and 10000 samples, the hierarchical selection is modestly less accurate than direct apricot but has performance close to that of direct apricot (Fig. 2). As *N* increases while keeping the same *rep set size*/*N*, the *d*_*HK*_ difference of the hierarchical selection minus direct apricot initially increases from negative values, then levels out, and then slightly decreases when *N* becomes very large, indicating that the performance difference between the hierarchical selection and direct apricot does not get larger when *N* further increases (Fig. 2). For the early and middle sets of samples, the trend of *d*_*HK*_ of the hierarchical selection is more similar to that of direct apricot, so their *d*_*HK*_ difference curves are flatter (Figs. 3 and 4). For the early sets, the *d*_*HK*_ difference between the hierarchical selection and direct apricot increases initially and then slightly decreases when *N* becomes very large (Fig. 3). For the middle sets, the *d*_*HK*_ difference still increases modestly when *N* becomes very large, however, the difference is still small (<0.081) (Fig. 4). Overall, these demonstrate that the hierarchical selection has performance close to that of direct apricot. Hierarchical selection’s close performance is partially contributed by the seeded-chunking and mean^2^-weighting.

The hierarchical selection substantially outperforms random sampling. Random sampling has substantially larger *d*_*HK*_ values than the hierarchical selection for all the recent, early, and middle sets of samples, and this trend is consistent (Figs. 2, 3, and 4). When the full set becomes very large, random sampling’s *d*_*HK*_ approaches 1.0 that is the maximum cosine distance. Random sampling follows the density distribution of the original set, so the rare cell/tissue types are not sufficiently represented, which yields large *d*_*HK*_.

### 3.2 Hierarchical representative set selection substantially reduces the run-time and memory usage of direct representative set selection, making selecting representative samples from the entire SRA set feasible

The hierarchical selection substantially reduces the runtime and memory usage of direct apricot. For the recent, early, and middle sets of samples, the hierarchical selection and direct apricot were all run using 85 cores for parallelism; their runtime and memory are reported in Tables S1, S2, and S3. As the full-set size increases, the real time and user+system time reductions increase (Tables 1, 2, and 3). When *N* = 10000, the user+system time reduction can reach 8.4X and the real time reduction can reach 7.47X. Real time reductions are less than user+system time reductions, since in direct apricot the full similarity matrix computation is fully parallel, while in the hierarchical selection although the similarity matrix computation for each chunk is fully parallel, chunks are processed sequentially to reduce the memory usage. Runtime reductions for most recent samples are generally greater than those for early- and mid-time samples, as longer reads generate more k-mers. The memory reduction also generally increases as the full-set size increases (Tables 1, 2, and 3). The memory reduction can reach 4.68X when *N* = 10000.

**Table 1.** Runtime and memory reduction with the hierarchical selection over direct apricot using the most recent 1000, 2000, 5000, 8000, 10000 samples in the SRA as the full sets.

| Reduction | Recent 1000<br>(select 100,<br>$l = 5$) | Recent 2000<br>(select 200,<br>$l = 10$) | Recent 5000<br>(select 500,<br>$l = 10$) | Recent 8000<br>(select 800,<br>$l = 10$) | Recent 10000<br>(select 1000,<br>$l = 10$) |
| --- | --- | --- | --- | --- | --- |
| Real time Reduction | 1.47 X | 2.72 X | 6.04 X | 6.53 X | 7.02 X |
| User+Sys time Reduction | 2.99 X | 5.76 X | 8.33 X | 8.17 X | 8.4 X |
| Memory Reduction | 2.13 X | 2.9 X | 4.47 X | 4.59 X | 4.54 X |

**Table 2.** Runtime and memory reduction with the hierarchical selection over direct apricot using the early-time 1000, 2000, 5000, 8000, 10000 samples in the SRA as the full sets.

| Reduction | Early 1000<br>(select 100,<br>$l = 5$) | Early 2000<br>(select 200,<br>$l = 10$) | Early 5000<br>(select 500,<br>$l = 10$) | Early 8000<br>(select 800,<br>$l = 10$) | Early 10000<br>(select 1000,<br>$l = 10$) |
| --- | --- | --- | --- | --- | --- |
| Real time Reduction | 1.53 X | 2.14 X | 4.62 X | 4.99 X | 7.47 X |
| User+Sys time Reduction | 3.05 X | 4.89 X | 5.88 X | 6.2 X | 8.31 X |
| Memory Reduction | 2.2 X | 2.83 X | 4.33 X | 4.37 X | 4.68 X |

**Table 3.** Runtime and memory reduction with the hierarchical selection over direct apricot using the mid-time 1000, 2000, 5000, 8000, 10000 samples in the SRA as the full sets.

| Reduction | Mid 1000<br>(select 100,<br>$l = 5$) | Mid 2000<br>(select 200,<br>$l = 10$) | Mid 5000<br>(select 500,<br>$l = 10$) | Mid 8000<br>(select 800,<br>$l = 10$) | Mid 10000<br>(select 1000,<br>$l = 10$) |
| --- | --- | --- | --- | --- | --- |
| Real time Reduction | 0.94 X | 2.24 X | 4.92 X | 5.66 X | 6.92 X |
| User+Sys time Reduction | 3.13 X | 5.5 X | 7.12 X | 7.1 X | 7.79 X |
| Memory Reduction | 2.33 X | 2.78 X | 4.58 X | 4.5 X | 4.55 X |

For the entire set of SRA RNA-seq samples (*N* = 196523), we use two levels of divide-and- merge (Fig. 1), with *m* = 1000, *Q* = *m*/5 = 200. At the 1st-level, *l* = 197, the random-subset-size *J* = 2000 (used for the seeded-chunking). At the 2nd-level, *l* = 40, *J* = 1000.

The hierarchical representative set selection makes selecting representative samples from the entire set of SRA RNA-seq samples feasible. When *N* = 10000, direct apricot by computing the full *N* × *N* similarity matrix takes 4.1 hours (using 85 cores) and 118.906 GB memory (Table S1). With the *O*(*N* ^2^) time complexity and *O*(*N*) space complexity, for the entire SRA set (*N* = 196523), the estimated runtime of direct apricot is 66 days (using 85 cores) and the estimated memory usage is 2336.776 GB, which is infeasible. The hierarchical selection on the entire SRA set (*N* = 196523) takes 17.6 hours (using 85 cores) and 56.430 GB memory, which makes this task fully feasible.

### 3.3 Hierarchical representative set selection outperforms random sampling for the entire set of SRA RNA-seq samples

The hierarchical representative set selection outperforms random sampling for the entire set of SRA RNA-seq samples. We compare the hierarchical selection with random sampling by selecting different sizes of representative sets from the entire SRA set (Fig. 5, Table S4). The hierarchical selection outperforms random selection in all these cases (Fig. 5); when selecting 7000 representative samples, the hierarchical selection outperforms random selection substantially, with a similar level of difference to those of smaller full sets. The *d*_*HK*_ values of the hierarchical selection are larger than that of selecting 1000 from the recent 10000 samples, mainly because the ratios *rep*_*set*_*size*/*full*_*set*_*size* are much smaller here, one magnitude smaller than the ratio 0.1 in selecting 1000 from 10000. As the size of the representative set increases, the performance of the hierarchical selection initially barely changes but increases when going from selecting 5000 to selecting 7000 representative samples. Random sampling’s *d*_*HK*_ are almost 1.0 (the maximum cosine distance) and do not change as the size of the representative set increases, indicating its poor performance on the entire SRA set.

**Figure 5.**
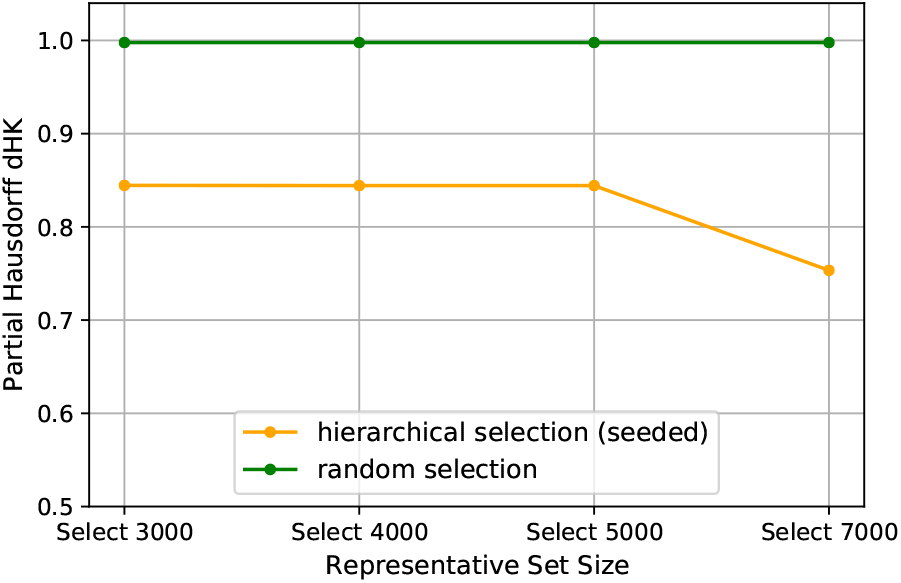
Selecting different sizes of representative sets from the SRA entire set (N =196523 human RNA-seq samples): partial Hausdorff distances d_HK_ of hierarchical selection and random selection. For d_HK_, q = 0.0001, so d_HK_ is the 21st-largest distance.

## 4 Discussion

Our results show that the hierarchical representative set selection is a close estimate to the direct representative set selection, while substantially reducing the runtime and memory usage of the direct selection, which makes subset selections feasible on big data such as the entire available RNA-seq samples in the SRA.

The chunk-size *m* needs to be large enough to avoid chunk overlaps in the seeded-chunking. If the chunk-size is too small, a large dense blob containing many more samples than the chunk-size could have multiple chunks overlapping there. Chunks need to have equal sizes (except for 1 or 2 chunks when *N*/*m* is not an integer) to fit to the resource capacity, so when the chunk of the closest seed is full, the data point has to be assigned to its closest non-full seed. So although the seeds are spread out, if the chunk-size is not large enough, there could still be chunk overlaps in the seeded-chunking.

The choices of the parameters *m* (which yields *l*), *Q*, and the number of iterations (levels) depend on the full-set size *N* and the memory of available computing resources. These parameters can affect the accuracy, runtime, and memory usage of the hierarchical selection. In all the results here, we use *Q* = *m*/5, which means *rep*_*set*_*size*/*chunk*_*size* = ∼0.2 for each chunk. Increasing this ratio could increase the selection accuracy at each chunk, however, the subsequent merged set would be larger, causing more chunks at the next level and thus increasing the runtime. Decreasing *m* (and thus increasing *l*) could reduce the runtime, however, smaller chunks have more overlaps which decrease the accuracy. Thus, the overall design of the hierarchy with parameters’ choices involves trade-offs between accuracy, runtime, and the resource capacity.

The mean^2^-weighting uses the average distance between samples of a chunk to indicate the chunk overall density. This is a heuristic. In a case that a chunk has several dense clusters that are far apart, causing a bigger average distance than the distances within clusters, the chunk may be assigned an unnecessarily larger weight. This may be partially addressed by performing clustering on each chunk and using the weighted-mean of average distances of all clusters as the chunk density. However, an accurate clustering incurs more computational cost. In our observation, most chunks do not have a distinctly clustered structure, and rather have a mixture of a few clusters and many roughly uniformly distributed data points. Thus, the mean^2^-weighting is a viable trade-off between runtime and accuracy.

In addition to the partial Hausdorff distance, a useful evaluation for a representative set could be using the representative set as the training set to train classifiers (that use the gene expression vectors as input features) to compare the accuracy. This is a direction for future work.

## 5 Conclusion

We demonstrate that our novel hierarchical representative set selection method can greatly reduce the runtime and memory usage of the direct representative set selection with the full similarity matrix computation, while still achieving performance close to that of the direct representative subset selection and substantially outperforming random sampling. This is the first approach to this problem that can scale to collections of the size of the full set of public human RNA-seq samples in the SRA.

## Acknowledgements

We would like to thank Dr. Guillaume Marçais for helping improve the speed of per-pair computation of k-mer similarity significantly. This work was supported by the National Science Foundation Graduate Research Fellowship Program [DGE1745016 to L.H.T.] (Any opinions, findings, and conclusions or recommendations expressed in this material are those of the authors and do not necessarily reflect the views of the National Science Foundation); the US NSF [grant 1937540 to C.K.]; the Gordon and Betty Moore Foundation’s Data-Driven Discovery Initiative [GBMF4554 to C.K.]; and the US National Institutes of Health [R01GM122935].

