## Supplementary Tables S1-S5 and Figures S1-S5 for "Practical selection of representative sets of RNA-seq samples using a hierarchical approach"

***Table S1*** *– Partial Hausdorff distance, classical Hausdorff distance, runtime, and memory usage of direct apricot, hierarchical selection, and random selection, using the most recent 1000, 2000, 5000, 8000, 10000 samples in the SRA as the full sets.*

| Set of samples | Method | Hausdorff $d_H$ | Partial Hausdorff $d_{HK}$ | Runtime (seconds) Real | User | Sys | Memory (GB) |
|---|---|---|---|---|---|---|---|
| Recent 1000 | direct apricot | 0.688321022 | 0.685257115 | 174.58 | 11070.16 | 111.42 | 17.131 |
| (select 100, | hierarchical (seeded) | 0.578381536 | 0.569809642 | 118.91 | 3449.28 | 292.08 | up to 8.061 |
| $l = 5$) | random selection | 0.837951394 | 0.826812255 | | — | | — |
| Recent 2000 | direct apricot | 0.664517495 | 0.662815581 | 655.18 | 42882.64 | 286.79 | 29.223 |
| (select 200, | hierarchical (seeded) | 0.650712434 | 0.637185175 | 240.77 | 6906.76 | 586.80 | up to 10.077 |
| $l = 10$) | random selection | 0.894947772 | 0.848467165 | | — | | — |
| Recent 5000 | direct apricot | 0.624796548 | 0.62353618 | 4334.92 | 306588.88 | 1323.23 | 67.515 |
| (select 500, | hierarchical (seeded) | 0.696114667 | 0.675768259 | 717.42 | 35803.41 | 1164.84 | up to 15.115 |
| $l = 10$) | random selection | 0.973292492 | 0.896965855 | | — | | — |
| Recent 8000 | direct apricot | 0.610919893 | 0.610735987 | 10095.85 | 701276.54 | 1513.85 | 101.776 |
| (select 800, | hierarchical (seeded) | 0.760030601 | 0.679049314 | 1545.26 | 84481.00 | 1588.09 | up to 22.169 |
| $l = 10$) | random selection | 0.991018754 | 0.904692245 | | — | | — |
| Recent 10000 | direct apricot | 0.607682609 | 0.607369442 | 14768.33 | 1047257.38 | 2273.86 | 118.906 |
| (select 1000, | hierarchical (seeded) | 0.70951932 | 0.665272021 | 2103.18 | 123182.85 | 1749.27 | up to 26.200 |
| $l = 10$) | random selection | 0.994547626 | 0.986377775 | | — | | — |

In Tables S1, S2, and S3, for $N$=1000, 2000, 5000, $d_{HK}$ is the 3rd largest distance; for $N$=8000, 10000, $d_{HK}$ is the 4th largest distance.

***Table S2*** – *Partial Hausdorff distance, classical Hausdorff distance, runtime, and memory usage of direct apricot, hierarchical selection, and random selection, using the early-time 1000, 2000, 5000, 8000, 10000 samples in the SRA as the full sets.*

| **Set of samples** | **Method** | **Hausdorff $d_H$** | **Partial Hausdorff $d_{HK}$** | **Runtime (seconds)** | | | **Memory (GB)** |
|---|---|---|---|---|---|---|---|
| | | | | **Real** | **User** | **Sys** | |
| Early 1000 | direct apricot | 0.725003411 | 0.723976221 | 153.22 | 7701.56 | 125.76 | 11.084 |
| (select 100, | hierarchical (seeded) | 0.745518016 | 0.725044029 | 100.12 | 2351.73 | 215.52 | up to 5.038 |
| $l = 5$) | random selection | 0.950139546 | 0.938242439 | | — | | — |
| Early 2000 | direct apricot | 0.67307199 | 0.672427473 | 401.49 | 22979.13 | 218.19 | 17.131 |
| (select 200, | hierarchical (seeded) | 0.760020571 | 0.706453874 | 187.64 | 4360.50 | 383.16 | up to 6.046 |
| $l = 10$) | random selection | 0.872261407 | 0.838447692 | | — | | — |
| Early 5000 | direct apricot | 0.61634404 | 0.615099659 | 2298.45 | 133723.63 | 565.12 | 39.300 |
| (select 500, | hierarchical (seeded) | 0.666160711 | 0.659981919 | 497.60 | 21561.05 | 1262.98 | up to 9.069 |
| $l = 10$) | random selection | 0.938632619 | 0.935902009 | | — | | — |
| Early 8000 | direct apricot | 0.630098995 | 0.628989605 | 5719.26 | 382221.70 | 874.59 | 70.538 |
| (select 800, | hierarchical (seeded) | 0.834517734 | 0.71208656 | 1145.94 | 60263.98 | 1509.99 | up to 16.123 |
| $l = 10$) | random selection | 0.9916713 | 0.908157434 | | — | | — |
| Early 10000 | direct apricot | 0.639209628 | 0.6383828 | 12315.44 | 800933.84 | 1446.46 | 89.684 |
| (select 1000, | hierarchical (seeded) | 0.837916798 | 0.713645489 | 1648.26 | 95117.21 | 1381.86 | up to 19.146 |
| $l = 10$) | random selection | 0.99772421 | 0.990244824 | | — | | — |

***Table S3*** – *Partial Hausdorff distance, classical Hausdorff distance, runtime, and memory usage of direct apricot, hierarchical selection, and random selection, using the mid-time 1000, 2000, 5000, 8000, 10000 samples in the SRA as the full sets.*

| **Set of samples** | **Method** | **Hausdorff $d_H$** | **Partial Hausdorff $d_{HK}$** | **Runtime (seconds)** | | | **Memory (GB)** |
|---|---|---|---|---|---|---|---|
| | | | | **Real** | **User** | **Sys** | |
| Mid 1000 | direct apricot | 0.666185918 | 0.664052819 | 150.21 | 9130.84 | 126.60 | 14.108 |
| (select 100, | hierarchical (seeded) | 0.685598613 | 0.648069325 | 160.45 | 2705.68 | 252.78 | up to 6.046 |
| $l = 5$) | random selection | 0.968587397 | 0.753610206 | | — | | — |
| Mid 2000 | direct apricot | 0.651423605 | 0.648054888 | 545.46 | 34587.00 | 248.80 | 25.192 |
| (select 200, | hierarchical (seeded) | 0.732927883 | 0.676261249 | 243.70 | 5827.03 | 508.06 | up to 9.069 |
| $l = 10$) | random selection | 0.967519071 | 0.923284417 | | — | | — |
| Mid 5000 | direct apricot | 0.646540816 | 0.646398158 | 3084.32 | 209037.81 | 892.53 | 55.422 |
| (select 500, | hierarchical (seeded) | 0.719052575 | 0.676710291 | 626.75 | 28509.77 | 983.47 | up to 12.092 |
| $l = 10$) | random selection | 0.961274547 | 0.867699133 | | — | | — |
| Mid 8000 | direct apricot | 0.636313583 | 0.635731787 | 7142.40 | 464515.34 | 865.10 | 81.622 |
| (select 800, | hierarchical (seeded) | 0.943464491 | 0.686230795 | 1262.73 | 64248.73 | 1301.38 | up to 18.138 |
| $l = 10$) | random selection | 0.998728362 | 0.989508253 | | — | | — |
| Mid 10000 | direct apricot | 0.654537313 | 0.653742747 | 12582.81 | 780770.97 | 1574.01 | 100.768 |
| (select 1000, | hierarchical (seeded) | 0.807148615 | 0.734005 | 1818.98 | 98855.02 | 1548.71 | up to 22.169 |
| $l = 10$) | random selection | 0.999886845 | 0.997812734 | | — | | — |

***Table S4*** – *Selecting different sizes of representative sets from the SRA entire set ($N$=196523 human RNA-seq samples): partial Hausdorff distance and classical Hausdorff distance of hierarchical selection and random selection.*

| **Metric** | **Select 3000** | | **Select 4000** | | **Select 5000** | | **Select 7000** | |
|---|---|---|---|---|---|---|---|---|
| | hierarchical (seeded) | random selection | hierarchical (seeded) | random selection | hierarchical (seeded) | random selection | hierarchical (seeded) | random selection |
| Hausdorff $d_H$ | 0.945978504 | 0.998763361 | 0.945978504 | 0.998763361 | 0.945978504 | 0.998763361 | 0.875009817 | 0.998763361 |
| Partial Hausdorff $d_{HK}$ | 0.844519904 | 0.997721045 | 0.844274154 | 0.997718649 | 0.844274154 | 0.997718649 | 0.753391502 | 0.997718289 |
| Representative -set-size/ Full-set-size | 0.0153 | | 0.0204 | | 0.0254 | | 0.0356 | |

***Table S5*** – *Performance comparison of hierarchical selection using seeded-chunking method vs. using sequential chunking method: partial Hausdorff distance and classical Hausdorff distance, using the most recent 1000, 2000, 5000, 8000, 10000 samples in the SRA as the full sets.*

| **Set of samples** | **Method** | **Hausdorff $d_H$** | **Partial Hausdorff $d_{HK}$** |
|---|---|---|---|
| Recent 1000 | hierarchical (seeded) | 0.578381536 | 0.569809642 |
| (select 100, $l = 5$) | hierarchical (sequential) | 0.612622717 | 0.578381536 |
| Recent 2000 | hierarchical (seeded) | 0.650712434 | 0.637185175 |
| (select 200, $l = 10$) | hierarchical (sequential) | 0.656343899 | 0.643755957 |
| Recent 5000 | hierarchical (seeded) | 0.696114667 | 0.675768259 |
| (select 500, $l = 10$) | hierarchical (sequential) | 0.72711968 | 0.708127649 |
| Recent 8000 | hierarchical (seeded) | 0.760030601 | 0.679049314 |
| (select 800, $l = 10$) | hierarchical (sequential) | 0.770009257 | 0.754661415 |
| Recent 10000 | hierarchical (seeded) | 0.70951932 | 0.665272021 |
| (select 1000, $l = 10$) | hierarchical (sequential) | 0.790246213 | 0.752879052 |

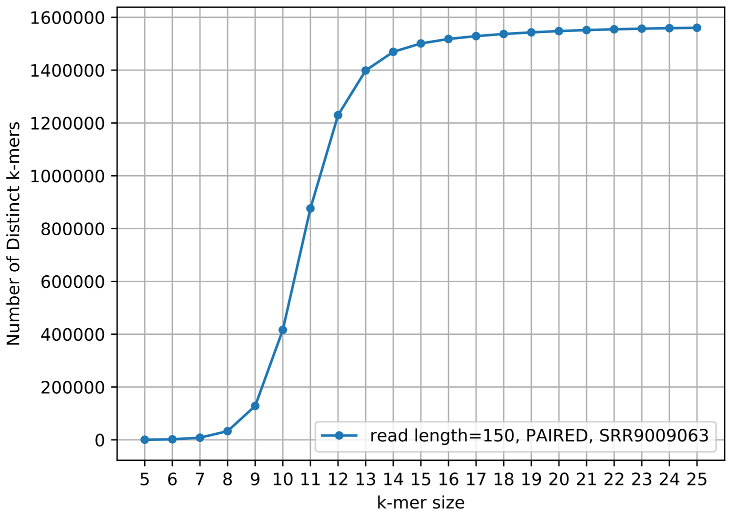


*Figure S1* – *The number of distinct k-mers vs. k-mer size: read-length=150, paired-end reads, from SRR9009063 (10000 random reads).*

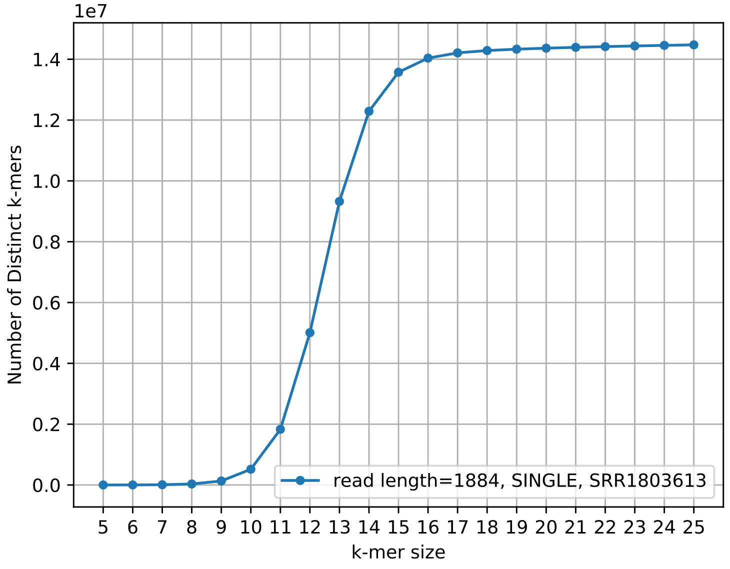


*Figure S2* – *The number of distinct k-mers vs. k-mer size: read-length=1884, single-end reads, from SRR1803613 (10000 random reads).*

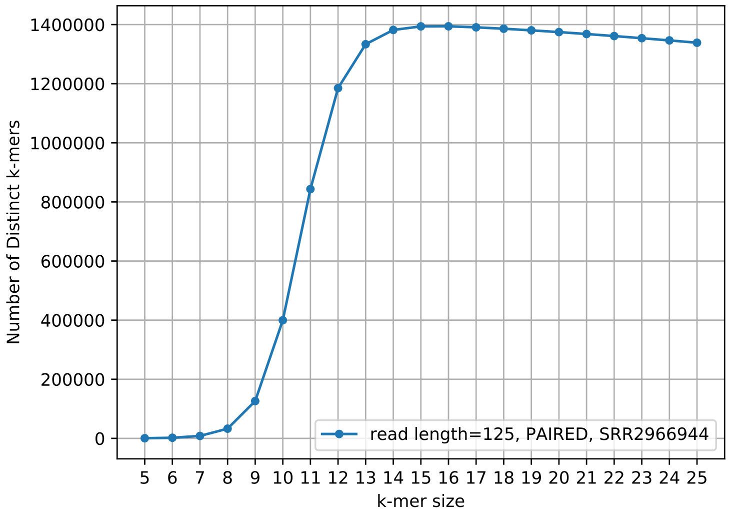


*Figure S3* – *The number of distinct k-mers vs. k-mer size: read-length=125, paired-end reads, from SRR2966944 (10000 random reads).*

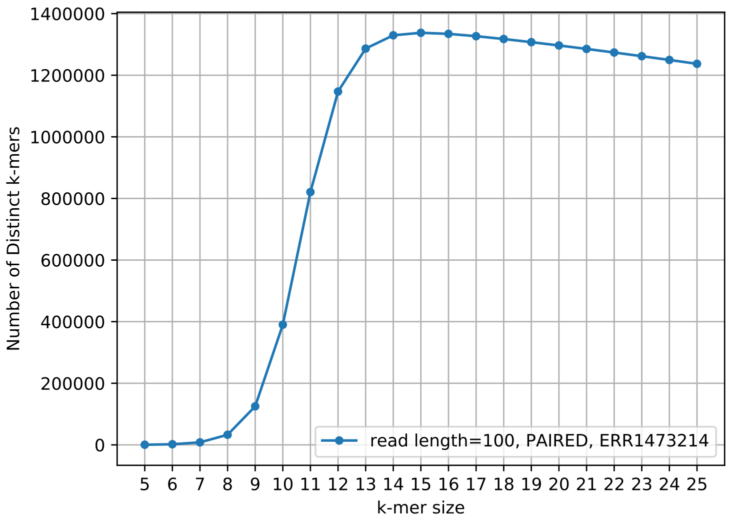


***Figure S4*** *– The number of distinct k-mers vs. k-mer size: read-length=100, paired-end reads, from ERR1473214 (10000 random reads).*

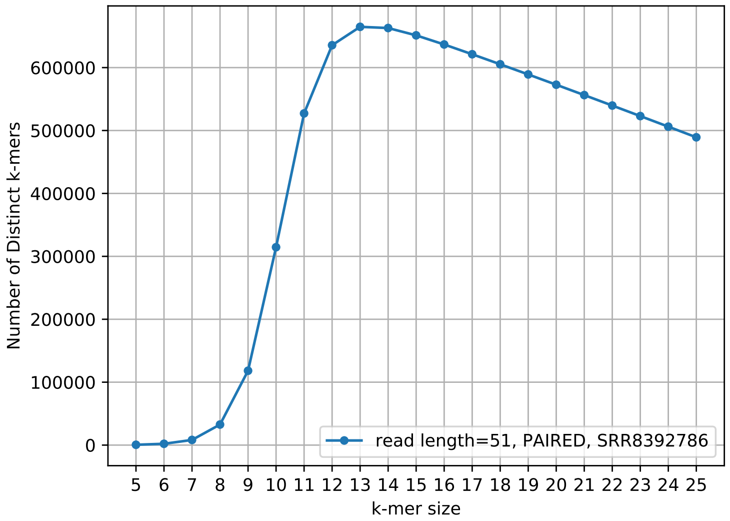


***Figure S5*** *– The number of distinct k-mers vs. k-mer size: read-length=51, paired-end reads, from SRR8392786 (10000 random reads).*

In Figs. S3, S4, and S5, the horizontal part of the curve bends down as k-mer size further increases, especially for shorter read-lengths, since when read-lengths are short, using larger k-mers would reduce the number of distinct k-mers.
